# Human microglia upregulate cytokine signatures and accelerate maturation of neural networks

**DOI:** 10.1101/2020.03.24.006874

**Authors:** Galina Schmunk, Chang N. Kim, Sarah S. Soliman, Matthew G. Keefe, Derek Bogdanoff, Dario Tejera, Ryan S. Ziffra, David Shin, Denise E. Allen, Bryant B. Chhun, Christopher S. McGinnis, Ethan A. Winkler, Adib A. Abla, Edward F. Chang, Zev J. Gartner, Shalin B. Mehta, Xianhua Piao, Keith B. Hengen, Tomasz J. Nowakowski

## Abstract

Microglia are the resident macrophages of the brain that emerge in early development and play vital role disease states, as well as in normal development. Many fundamental questions about microglia diversity and function during human brain development remain unanswered, as we currently lack cellular-resolution datasets focusing on microglia in developing primary tissue, or experimental strategies for interrogating their function. Here, we report an integrative analysis of microglia throughout human brain development, which reveals molecular signatures of stepwise maturation, as well as human-specific cytokine-associated subtype that emerges around the onset of neurogenesis. To demonstrate the utility of this atlas, we have compared microglia across several culture models, including cultured primary microglia, pluripotent stem cell-derived microglia. We identify gene expression signatures differentially recruited and attenuated across experimental models, which will accelerate functional characterization of microglia across perturbations, species, and disease conditions. Finally, we identify a role for human microglia in development of synchronized network activity using a xenotransplantation model of human microglia into cerebral organoids.

## Introduction

As key resident macrophages of the brain, microglia play vital roles in responding to disease states, such as viral infections, injury, or brain tumors (*1–5*). Mutations in microglia-related genes have been implicated in a wide range of neurological (*6, 7*), neuropsychiatric (*8, 9*), and neurodegenerative diseases (*10*). As part of their response to disease environments, microglia have been shown to adopt novel transcriptional states (*11–13*), therefore transcriptomic characterization of microglia heterogeneity could reveal pathways that could be harnessed to control the immune response in disease states.

Unbiased high-throughput single cell transcriptomics (scRNAseq) of adult microglia has recently revealed deep evolutionary conservation of the core molecular signature, as well as striking species specific variations in metabolic and immune genes (*14, 15*). However, resting adult microglia are molecularly homogeneous, with relatively subtle differences in gene expression characterizing microglia across anatomically defined brain regions (*16*). In contrast, microglia of the developing brain are highly heterogeneous and follow stepwise maturation trajectory (*17*). Whether human microglia undergo similar transitions is currently unclear (*18*), and comprehensive characterization of prenatal human microglia is needed to benchmark and improve *in vitro* differentiation protocols (*19–21*).

To define molecular subtypes of microglia during human brain development, we *in silico* sorted cells that expressed high levels of previously published microglia markers (*22*) from existing BRAIN Initiative Cell Census Network (BICCN) datasets (Fig. 1A, fig. S1)(see Materials and Methods). We identified 20,315 putative microglia cells (table S1) derived from four anatomically-defined regions (cerebral cortex, ganglionic eminence, hypothalamus, thalamus) across a range of developmental periods between gestational week (GW) 4 and 25 that passed basic quality control metrics (fig. S1-S2). In addition, we incorporated recently published data from the human hippocampus (*23*). We corrected batch effects (fig. S1C) using Harmony (fig. S1G)(*24*), and embedded single cell data in nearest neighbor space (Fig. 1B). We detected eight molecularly distinct cell types using Louvain clustering (Fig. 1B) and identified marker genes for each cluster using a general linear model (Fig. 1C, fig S1-S2, table S2). Two clusters expressed markers of lymphoid lineage consistent with B cells (CD79B, major histocompatibility complex II genes MHC-II) and natural killer cells (NKG7, KLRB1), and one cluster expressed genes involved in erythropoiesis (HBG2, HBA2, HBB) that may represent early progenitors of erythromyeloid lineage (fig. S1I). We removed these three populations from further analysis. Brain region and sample age were identified as major sources of transcriptional variance (Fig. 1D-E, fig. S1). All clusters were composed of cells from multiple individuals and brain regions, suggesting that molecularly distinct microglia subtypes can be found broadly across the developing telencephalon and diencephalon (fig. S1). We validated the spatial distribution of the major clusters using single molecule fluorescence *in situ* hybridization (smFISH) for key markers of individual clusters across two specimens around midgestation (Fig. 1F-G, fig. S2C).

**Fig. 1.**
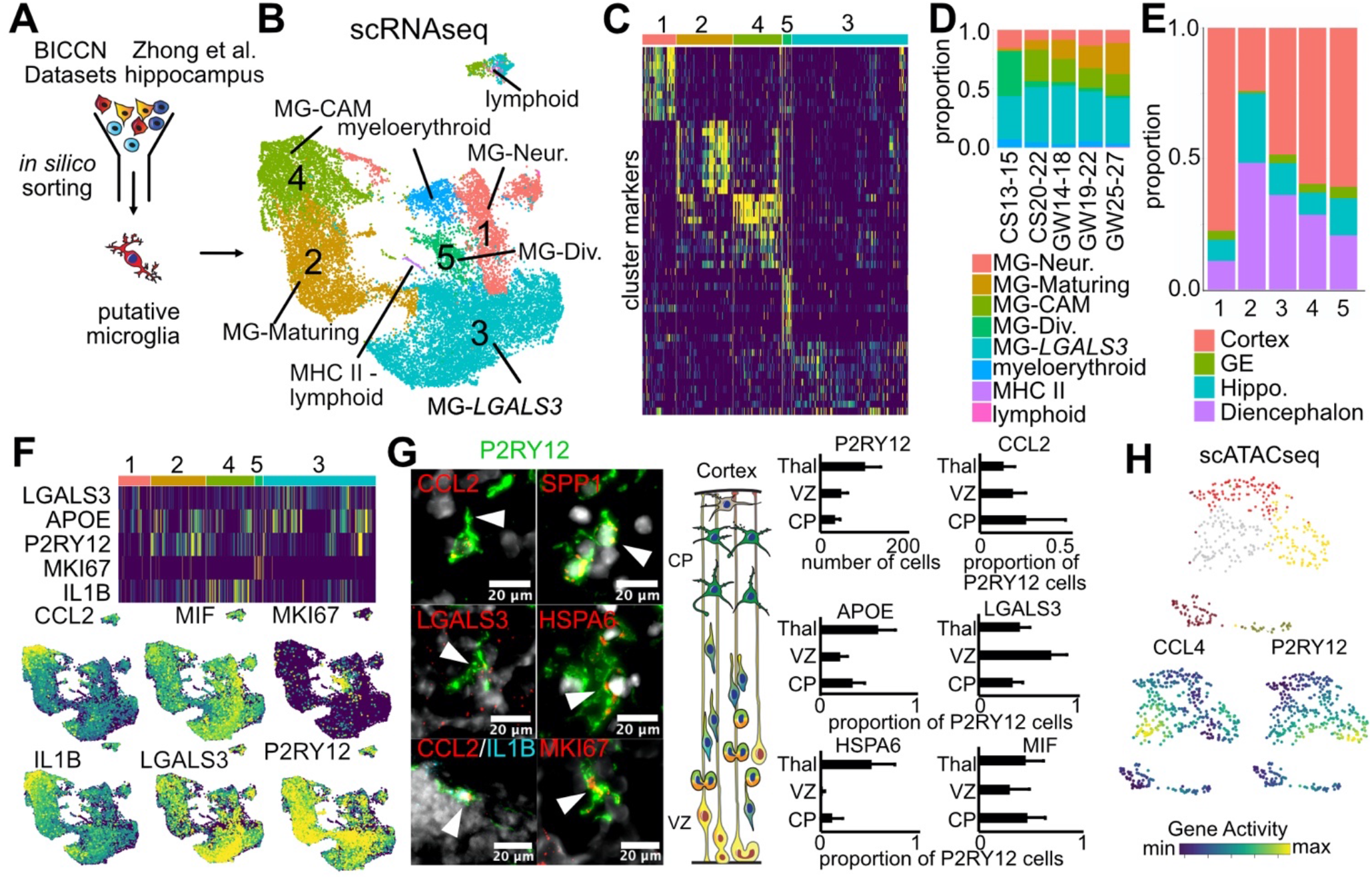
Molecularly distinct subtypes of microglia are present in the developing human brain. **(A)** Primary human microglia from GW6-GW25 were *in silico* purified from the BICNN dataset (total of 17,444 cells) and combined with microglia cells from hippocampus (total of 2,871 cells from (20)). **(B)** UMAP plot of cells passing selection criteria. Unbiased clustering of 20,315 cells from 33 individuals identifies 5 clusters of microglia and three non-microglial cell types. Lymphoid cells robustly express markers of NK cells (NKG7 and KLRB1), and MHC II lymphoid cluster is enriched for B cell markers (CD79B and MHC II genes). **(C)** Heatmap of top cluster-enriched genes across five microglia clusters. **(D)** Cluster proportion for each age range. CS, Carnegie stage (CS13-15 = GW6-7; CS20-22 = GW10); GW, gestational week **(E)** Brain region compositions of each cluster; GE, ganglionic eminence; hippo, hippocampus **(F)** Heatmap and feature plots of key cluster-enriched genes that were selected for smFISH validation. **(G)** smFISH validation of top cluster marker expression in human tissue. Thal – thalamus; VZ- ventricular zone; CP – cortical plate. **(H)** Single cell chromatin state profiling (scATACseq) identifies molecular subtypes of microglia, including distinction between cells with enriched accessibility of *CCL4,* equivalent to MG-CAM from **(B)** and *P2RY12*-enriched cluster equivalent of cluster 2 in **(B)**.

Among five high-confidence microglia clusters, Cluster 1 was enriched for cells captured from the cerebral cortex and showed enrichment for neuronal genes, such as *SOX4, STMN2*, and *TUBA1A*, and we refer to it as “MG-Neur.” (Fig. 1A-E, fig. S1H-J, fig. S2B). These cells also expressed elevated levels of some of the autism-relevant genes compared with other microglia clusters (fig. S3). Expression of neuronal genes could indicate, consistent with previous reports, that this cluster represents microglia involved in active phagocytosis of newborn neurons (*25*). Consistent with this hypothesis, cluster 1 was enriched for cells sampled from germinal zones (fig. S1I-J), but we did not see any substantial enrichment of phagocytosis-related genes (fig. S2F).

Cells in Cluster 2 expressed genes related to heat shock response (*HSPA6, HSPA1B, HSPB1*), and many immediate early genes (*JUN* and *FOS*) (Fig. 1C, fig. S1G). This gene expression signature was recently reported in adult microglia (*17*), but could represent an artifact of cell isolation. Consistently, we did not detect substantial levels of *HSPA6* mRNA in cerebral cortex tissue sections (Fig. 1G). However, this cluster was also enriched classical markers of mature microglia: *P2RY12, SALL1, CSF1R,* and *CX3CR1*, its relative abundance increased with sample age (Fig. 1D), suggesting that Cluster 2 could represent more mature microglia consistent with a recent report (*17*). Interestingly, this cluster was enriched for cells sampled from hippocampus and diencephalon (hypothalamus and thalamus) (Fig. 1E-G, fig S1F), suggesting that while microglia follow a stepwise maturation process as a population, the rates of maturation may differ across brain regions.

The largest cluster of microglia (cluster 3) was enriched for *APOC1, FTL, S100A11*, and *LGALS3* (Fig. 1C, 1F, fig. S1H, J, S2B). This expression pattern has been recently reported in early postnatal mouse brain, and shown to be associated with proliferative zones (*13*) and white matter (*4, 26, 27*). This cluster was found across all stages of development and all brain regions (Fig 1D-E).

We also identified cluster 4 that was enriched for genes involved in cytokine signaling, *CCL2*, *CCL4*, *CCL4L2*, *TNF* and *IL1B* (*4, 28*), and human immunodeficiency virus entry cofactor *CXCR4 (29)* (Fig. 1B-G,figs. S1 and S2, table S2). Interestingly, a similar signature was recently shown to be human specific based on a single cell RNA seq study from the adult brain (*14*). Given the short processing time of our sample, we predict that these cytokine-associated microglia (CAM) represent a *bona fide* subtype of microglia present *in vivo. CCL2* was predominantly expressed in microglia from the cortical plate (Fig. 1G, fig. S1I). What could underlie the evolutionary changes in gene expression is unclear, but interestingly, *CCL4* and *CCL4L2* coding loci are part of a recent segmental duplication on chromosome 17 that also created an extra copy of *TBC1D3*, recently implicated in evolutionary expansion of the cerebral cortex (*30*). It is possible that genomic changes in the nearby noncoding DNA contributed to the expanded expression of *CCL4* in the human microglia compared to other species. Species specific differences in gene expression could underlie changes in morphology and cell dynamics across species (*14*). For example, *CCL3* (MIP-1alpha) and *CCL2* (MCP-1) regulate microglia morphology and motility under normal conditions, but were previously thought to be astrocyte-derived (*31*).

To further explore the molecular signatures of prenatal microglia, we analyzed chromatin accessibility data generated using single cell ATAC-seq (*32*), and we further annotated putative enhancer elements among open chromatin regions by performing genomic profiling of H3K27ac CUT&TAG (*33*). Clustering of microglia based on scATACseq profiles revealed strong separation of epigenetic profiles between *CCL4*-positive and *P2RY12*-positive microglia clusters, mirroring the distinction from scRNAseq (Fig. 1H, fig. S2E). To predict what transcription factors motifs are enriched in either cluster, we identified cluster-specific ATAC-seq peaks using Fisher’s exact test and predicted transcription factor motif enrichments using HOMER (*34*). Transcription factors related to cytokine signaling and immediate early gene expression were significantly enriched in the cytokine associated cluster (Fig. S2D). Finally, we predicted enhancer-gene interactions using the activity-by-contact algorithm (*35*), which identified 48,819 predicted enhancers that interact with genes (table S3).

Finally, we also identified a cluster of microglia progenitors (cluster 5) that expressed high levels of cell cycle genes. Although dividing microglia represent only a modest fraction of all cells analyzed, this cluster was enriched for first trimester cells (Fig. 1D, fig. S1E). Microglia precursors colonize the developing brain starting around gestational week GW6 and are derived from the yolk sac (*36, 37*). Primitive microglia progenitors are first located in the subplate and subventricular zone before expanding into other brain regions (fig. S1K-L)(*25, 38, 39*). Iterative analysis of cells derived from first trimester samples revealed a molecularly distinct population of *MS4A7* monocyte-like cells (*17*) (Fig.2A-C, fig S4A-D). Consistent with a previous report in mice (*26*), first trimester microglia expressed high levels of *F13A1, LYVE1, MRC1* and *DAB2.* Starting at around GW8, expression of more canonical microglia marker genes *CX3CR1, C3, SALL1*, and *TREM2* markedly increased (Fig. 2C).

**Fig. 2.**
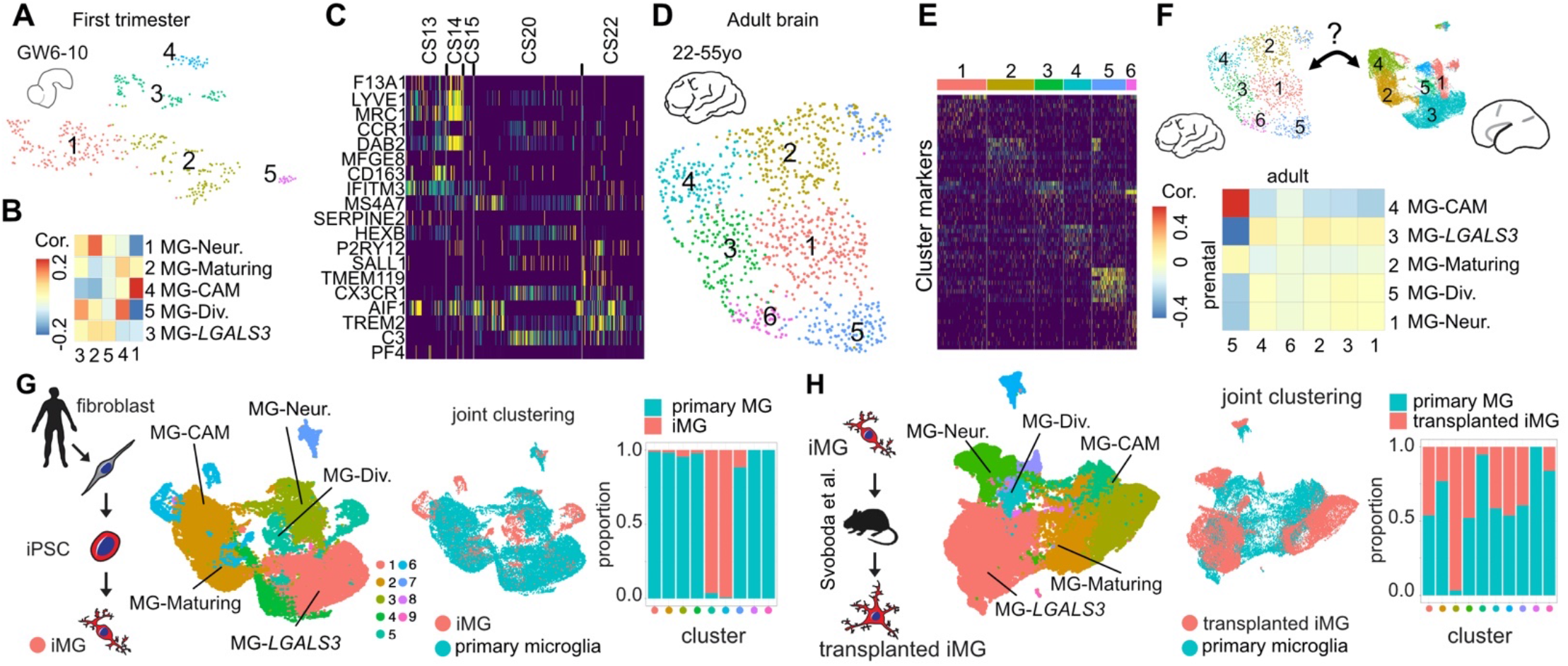
Microglia cluster signatures and composition differ across timepoints and experimental models. **(A)** UMAP of clusters of first trimester human microglia. 436 cells from six individuals formed five clusters. **(B)** Cluster correlation between first and second trimester microglia demonstrates high correlation for CAMs. **(C)** Heatmap of macrophage and microglia markers demonstrate transition from macrophage-like signature to microglia markers between CS13 and 22 (GW6-10). **(D)** UMAP of five clusters of adult microglia. **(E)** Heatmap of top ten differentially expressed genes for each cluster across adult microglia. **(F)** Inter-cluster correlation between adult and primary prenatal clusters identifies CAMs as the most preserved microglia subtype between prenatal and adult microglia. **(G)** Joint analysis of 20,315 primary and 2,838 induced microglia (iMG) cells. ìMG forms several individual clusters that partially overlap with primary prenatal microglia cells. **(H)** Joint analysis of primary microglia with 17,151 cells from induced microglia transplanted into mouse brain (from (*19*)). Individual clusters are comprised of both cells from *in vivo*-transplanted iMG and primary microglia.

Together, this cellular resolution atlas of human microglia provides a reference for comparing microglia signatures across developmental stages and experimental models. To reconstruct the developmental trajectory between developing and adult human microglia, we compared prenatal scRNAseq data to the recently published scRNAseq microglia dataset from the adult brain (22-55 year old samples) generated using MARS-seq (Fig. 2D-E, fig. S4E)(*14*). Unbiased analysis and pairwise cluster correlation analysis revealed *CCL4*-enriched cluster as the most conserved signature between development and adulthood (Fig. 2E-F, fig. S4E). Signatures associated with *LGALS3*-enriched cluster were not detected in adult microglia, consistent with previous reports that these clusters are associated with early postnatal mouse microglia (*13, 26*), while markers of maturing microglia, such as *TMEM119, P2RY12, CX3CR1* that are enriched in developing microglia Cluster 2 could be found broadly expressed across all adult microglia clusters (fig. S4E). Together, this analysis further emphasizes the remarkable conservation of cytokine associated microglia across datasets, experimental platforms, and developmental stages.

Next, we sought to leverage developing brain microglia datasets to benchmark common experimental models of this cell type. Efficient culture models of human microglia are required to advance our understanding of normal brain development and neuroimmune interactions in disease states. However, the dynamic nature of microglia gene expression and morphology (*40*) represents a significant challenge for studies involving human microglia. We generated scRNAseq data for microglia that were induced from pluripotent stem cells using a recently established protocol (*21*) (see Materials and Methods), and we analyzed recently published data for induced microglia that had been transplanted into *in vivo* mouse brain (*19*). We co-embedded both datasets in the space of primary microglia and compared the level of mixing with primary cell clusters (Fig. 2 G-H). Remarkably, while iMG cells clustered away from primary cell clusters, they developed some of the key subtype-specific signatures, including the cytokine signature, and subtype enriched for *LGALS3* (Fig. 2G, fig. S4F-N). In striking contrast, microglia that were transplanted into the mouse brain recovered many of the primary cell signatures (Fig. 2H, fig. S4F-N), as has been previously reported (*19*), with the exception of cytokine-associated microglia that formed two separate clusters composed of primary and transplanted cells, respectively (Fig. 2H, fig. S4H,K). Together, this analysis revealed that many important features of microglia diversity are recapitulated in an *in vitro* model and are further improved by xenotransplantation into the mouse brain, with the exception of the cytokine associated microglia that remain distinct from their primary counterparts.

Subsequently, we sought to compare *in vitro* models of primary human microglia. To isolate microglia from primary human tissue, we isolated microglia from primary human tissue using magnetic beads for CD11b (see Materials and Methods) and performed scRNAseq to identify what microglia subtypes can be isolated using this strategy. We were able to recover all of the major microglia subtypes, including *LGALS3*-positive, cytokine-associated, and dividing subtypes (Fig. 3B-E, fig S5). We noted distinct clusters of macrophages (*LYVE1/F13A1*-positive), putative endothelial cells (*CLDN5/ITM2A*-positive). After 8 days of *in vitro* culture (*41*), we recovered many of the same populations of microglia as the original dataset, with the exception that *in vitro* cultured microglia appear to downregulate immediate early genes (*JUN, RHOB, HSPA1A, DNAJB1*), but upregulated neuronal marker genes found in cluster 1 of acutely processed primary microglia (*SOX4, STMN2, SOX11*) which appears to be lost during microglia purification (fig. S6). Although cultured primary microglia expressed lower levels of *P2RY12* mRNA (fig. S7A), which is downregulated *in vitro (40, 41),* most cells expressed detectable levels of P2RY12 protein (Fig. 3F), and were amenable to lentiviral transfection of reporter gene (Fig S7D). Interestingly, mouse microglia cultured under the same conditions, had larger cell body size and were more ramified (Fig. S7B). These results suggest that differences between microglia across species are not limited to molecular signatures, but are also reflected in cell-intrinsic morphology and dynamic cell behavior.

**Fig. 3.**
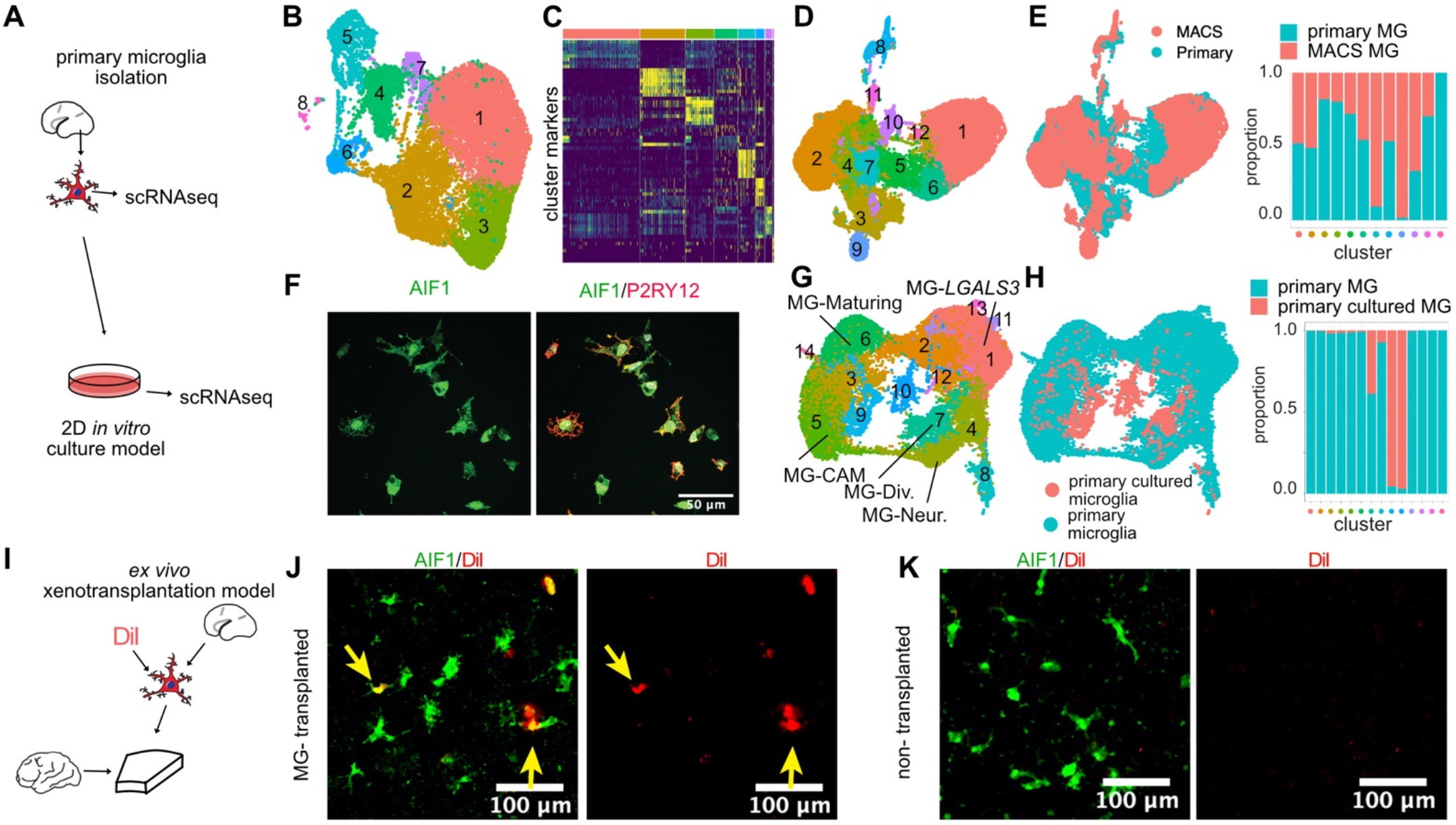
Experimental models for microglia culture and xenotransplantation into brain tissue. **(A)** Primary prenatal microglia were purified with MACS-sorting and used for scRNAseq before and after *in vitro* culture. **(B)** UMAP of 16,876 MACS-purified microglia from GW23 that were acutely sequenced right after purification. **(C)** Clustering of microglia reveals molecularly heterogeneous subtypes. **(D)** Joint embedding and clustering of MACS-purified microglia and primary *in silico*-sorted microglia form nine unique clusters **(E)** Metadata annotation of the co-embedded object. Co-clustered MACS-purified and primary prenatal microglia demonstrate extensive overlap across all microglia subtypes in both samples. Bar plot represents the relative contribution of purified and acutely profiled microglia across clusters. **(F)** Purified primary human microglia can be cultured in two dimensions and expresses P2RY12 *in vitro*. **(G)** UMAP of 1,414 cultured primary microglia co-embedded with primary non-purified cells and clustered. **(H)** Relative contribution of cultured and acutely profiled microglia across clusters. **(J)** Xenotransplantation model of human microglia into adult brain tissue slice cultured *ex vivo*. Adult human slices were maintained in culture for two days and then MACS-purified microglia labeled with DiI were added to the slices. **(K)** Transplanted microglia (labelled with DiI) after 5 days in culture integrate into brain tissue. **(L)** Control nontransplanted tissue slices maintain microglia in culture.

The extreme plasticity and migratory capacity of microglia makes them a good candidate for transplantation therapies, and several studies have proposed microglia transplantation as a therapeutic avenue for Alzheimer’s disease (*42*) and neurodevelopmental diseases (*43, 44*). Given the extreme differences in microglia transcriptomics across species, as well as significant plasticity of microglia across environmental conditions, we reasoned that additional experimental models of human tissue cultures could represent an important intermediate for understanding the barriers to human brain transplantation of microglia. We sought to explore the feasibility of xenotransplantation of human microglia into human organotypic slice cultured *ex vivo.* Traditional organotypic slice culture methods often are not permissive to maintaining microglia for protracted time periods. We first compared several of the major slice culture media protocols and found that serum-free media supplemented with IL34, TGF beta and cholesterol conditions allow us to maintain microglia in cultured organotypic brain tissue slice for at least a week (fig. S7F-H).

Subsequently, we prepared brain tissue slice culture from adult human neurosurgical resection and performed transplantation of primary prenatal microglia that were labeled with a lipophilic dye DiI. After five days in culture, transplanted microglia were located throughout the recipient slice, maintained ramified morphology, expressed AIF1 and P2RY12 (Fig.3I-K), and were highly motile (Movie S1). These experiments establish proof of concept that human microglia could be transplanted into adult tissue, and this experimental model could provide a tractable experimental system for exploring barriers to efficient cell transplantation.

Addressing long-standing questions about the role of microglia in the early stages of human neurogenesis and neuronal maturation is challenging due to the lack of experimental models. Microglia have been shown to interact with neural stem cells and newborn neurons (*25, 39, 45*), but the normal physiological roles are poorly understood. *In vitro* differentiation protocols of human pluripotent stem cells are becoming widely adopted to derive cultured models of developing brain (*46*). Unlike neurons, astrocytes, and oligodendrocytes, mature microglia do not emerge in organoids in high numbers (*47*), but can be reconstituted by transplantation (*47*). This strategy offers an opportunity to dissect the roles of microglia in the normal developing brain. We differentiated human brain organoids and at 5 weeks of differentiation, we transplanted primary MACS-purified human microglia from midgestation specimens (Fig. 4A, Movies S2 and S3). Over the course of 5 days, microglia invaded the organoid, and transformed their morphology from migratory to ramified state and can be detected even 5 weeks after transplantation (Fig. 4B, fig. S8). Thus, the ‘neuroimmune’ organoid offers an opportunity to explore the long-term impact of microglia on human neurogenesis and neuronal maturation. Towards this goal, we profiled neuroimmune organoids generated from three different iPS lines at 5 weeks after transplantation using scRNAseq. We identified the major cell types, including cells of forebrain and hindbrain identity (Fig. S8A-C). Off-target lineages can emerge with different efficiencies across the three iPS lines used, as we previously reported (*48*). After selecting for cells of forebrain (*FOXG1*-positive) identity, iterative clustering revealed the major cell types of the excitatory neuronal lineage, radial glia, intermediate progenitors, and neurons (Fig. 4C). Cells derived from control and neuroimmune organoids contributed across all clusters. Differential gene expression analysis revealed no genes differentially expressed between neuroimmune and control cortical neurons, but many genes differentially expressed at the level of radial glia and intermediate progenitors (Fig. 4E, fig. S8F-G), including significant reduction of genes involved in apoptotic cell death among ‘neuroimmune’ radial glia. Interestingly, we found that the number of differentially expressed genes decreased along the differentiation axis, suggesting that radial glia and neural progenitors may be more sensitive to the presence of microglia than neurons at the transcriptional level. Genes that showed expression bias towards higher levels in neuroimmune organoids included markers of synaptic maturation, *MEF2C* and *NRXN1*. On the other hand, markers of neurogenesis and immature neurons (*NEUROD2, TUBB, ID2*) showed bias towards higher expression in control organoids (fig. S8H-I). Although these expression differences did not meet p-value criteria for differential expression, we decided to explore whether microglia might contribute to neuronal maturation using orthogonal approaches. First, immunostained organoid sections for vGlut1 and PSD95 revealed significant reduction in synaptic puncta (Fig. 4F), consistent with the role of microglia in synaptic remodeling (*49, 50*). In addition, we explored whether presence of microglia might influence functional network maturation that is known to occur in organoids (*51*). We recorded spontaneous neural activity using multi-electrode arrays in neuroimmune and control organoids at 5 weeks after transplantation and found increased synchronization and frequency of oscillatory bursts in neuroimmune organoids compared to control (Fig. 4G, fig. S9), consistent with accelerated network maturation in organoids (*51*) and with the presumed role of microglia in network formation (*52, 53*). These findings suggest that microglia transplant into organoids long-term and influence brain development via their modulation of spontaneous oscillatory network properties through their interaction with developing synapses.

In summary, our analysis of transcriptional and epigenetic heterogeneity of microglia in the developing human brain highlights dynamic developmental transitions that mimic stepwise maturation of mouse microglia. We report that cytokine-associated microglia expressing elevated levels of CCL2, CCL4, and IL1B, recently suggested as human-specific, are present in early brain development. In addition, we characterized and compared strategies for microglia isolation and culture that could serve as experimental systems for functional characterization of human microglia, including two-dimensional culture and transplantation into cerebral organoids. We examined the role of microglia in the development of human neural circuits, and demonstrated an impact of transplanted microglia on network-level synchronized activity characteristic of developing cortical networks in cerebral organoids.

**Fig. 4.**
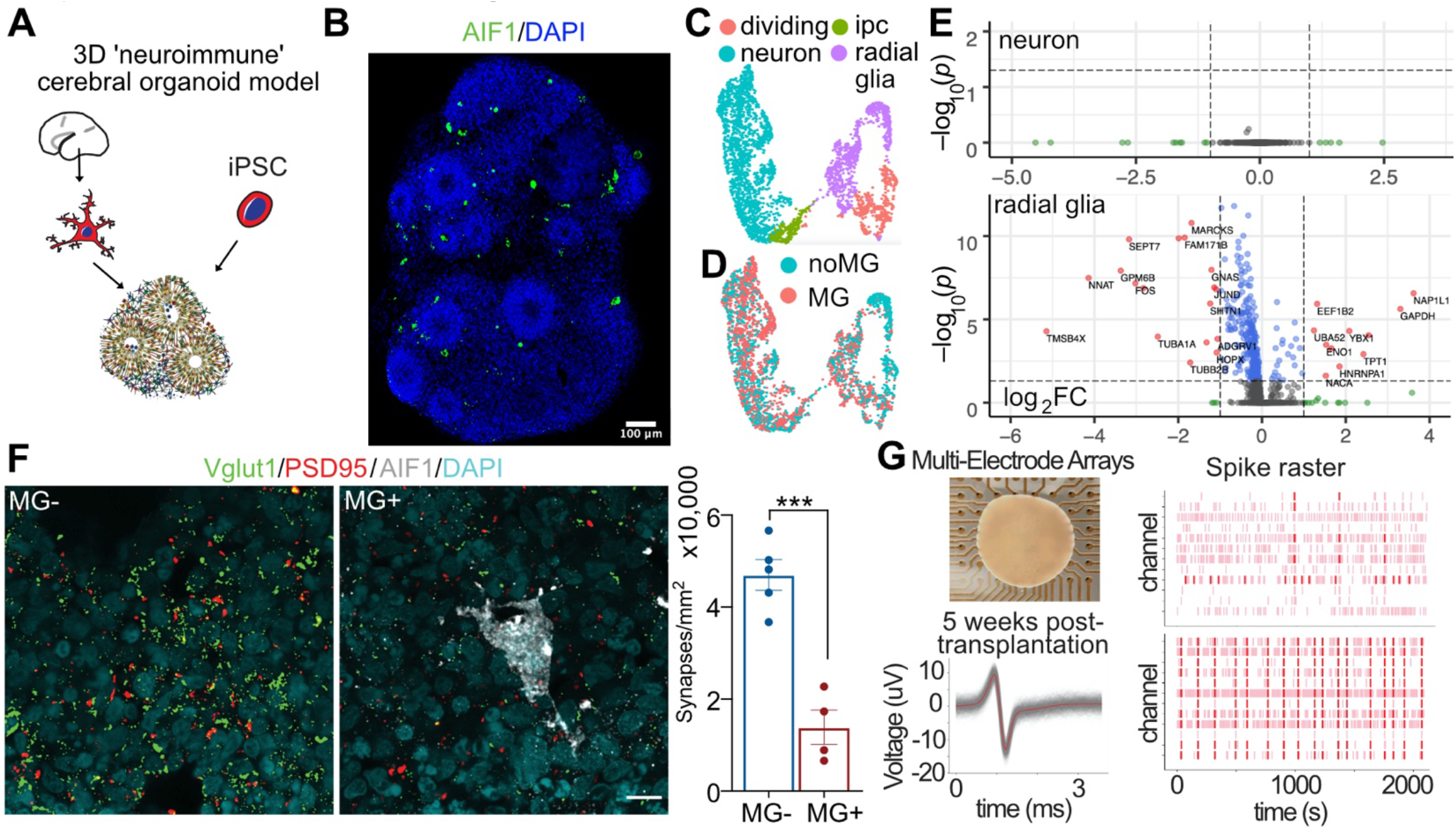
Chimeric cerebral organoid model for studying neuroimmune interactions during human brain development. **(A)** Generation of ‘neuroimmune’ organoids through xenotransplantation of primary prenatal microglia into cerebral organoids.**(B)** Microglia are present within cerebral organoid 5 weeks after transplantation. **(C)** UMAP of scRNAseq data of FOXG1-positive organoid cells. **(D)** Cells from control and ‘neuroimmune’ organoids contribute across all clusters. **(E)** Differential gene expression analysis reveals robust transcriptomic differences in radial glia but not neurons. **(F)** Microglia transplanted organoids show reduced density of synapses. **(G)** Microglia transplantation promotes synchronized burst activity in cerebral organoids.

## ACKNOWLEDGMENTS

We thank Drs. Aparna Bhaduri and Arnold Kriegstein for generously facilitating the access to BICCN datasets. We thank all members of the Nowakowski, Mehta and Piao laboratories for helpful discussions and comments throughout this project.

## Materials and Methods

### Consent statement UCSF

De-identified tissue samples were collected with previous patient consent in strict observance of the legal and institutional ethical regulations. Protocols were approved by the Human Gamete, Embryo, and Stem Cell Research Committee (institutional review board) at the University of California, San Francisco.

### Single cell RNA sequencing

Single cell RNA-seq libraries were generated using the 10x Genomics Chromium 3’ Gene Expression Kit. Briefly, single cells were loaded onto chromium chips with a capture target of 10,000 cells per sample. Libraries were prepared following the provided protocol and sequenced on an Illumina NovaSeq with a targeted sequencing depth of 50,000 reads per cell. BCL files from sequencing were then used as inputs to the 10X Genomics Cell Ranger pipeline.

### Single cell RNA-seq Analysis

For preprocessing of scRNA-seq data, CellRanger was used to create a cell by gene matrix which was then processed using Cellbender (*1*) to remove ambient RNA from every run and Solo (*2*) for doublet detection and removal. A minimum of 200 genes, 500 UMI counts, and 20% mitochondrial cutoff were used to remove low quality cells from all datasets. To *in silico* sort microglia from the BICCN datasets, we selected cells expressing 10 or more cumulative UMI counts of the following genes: CCL3L3, CCL4, C3, CCL3, PLEK, FOLR2, ITGAX, SPP1, CSF3R, BIN2, DHRS9, CD74, CD69, IL1B, CX3CR1, OLR1, CH25H, FCGR2A, ADORA3, LAPTM5, P2RY12, IRF8, AIF1. The dataset was further filtered for non-microglial cells after dimensional reduction based on preliminary clustering and marker genes that were not exhibiting canonical signatures, or other cell types. The SCTransform (*3*) workflow was used and then PCA was computed on the residuals for input into Harmony (*4*) for batch correction. The parameters of Harmony were set to use the top 10 principal components with theta set to 20. Uniform manifold approximation and project (UMAP) (*5*) embeddings and neighbors for Leiden clustering (*6*) used the 10 components outputted from Harmony. Organoid demultiplexing and doublet filtering was done through deMULTIplex (https://github.com/chris-mcginnis-ucsf/MULTI-seq). Pearson correlation was calculated on the intersection of the shared genes between datasets which averaged Pearson residuals for each cluster.

### Differential Expression

MAST (*7*) was used on log normalized raw counts for all differential expression tests, except for the organoids. Wilcoxon ranked sum test was used on Pearson residuals from SCTransform for increased sensitivity on microglia treated organoids. Volcano plots were set to have a threshold of Bonferroni corrected p-value of 0.05.

### Single cell ATAC-seq analysis

scATAC-seq data were generated and analyzed as previously described (*8*). Briefly, nuclei were isolated from mid-gestation human cortical samples and scATAC-seq libraries were generated using the 10X Genomics scATAC-seq solution. Sequencing libraries were processed using the snapATAC pipeline (*9*). Batches were integrated using a custom implementation of scAlign (*10*). After cell types were identified using a classifier model based on scRNA-seq data from similar samples, microglia cells were isolated for downstream analysis. Dimensionality reduction, clustering, and gene activity scoring were performed as previously described (*8*). Peaks were called using MACS2 on the combined signal from all microglia cells. Cluster-specific peaks were identified using a Fisher’s Exact test and a p-value cut-off of 0.05. The findMotifsGenome.pl function from the HOMER package was used for identifying transcription factor motif enrichments in differentially accessible peaks.

### CUT&Tag data generation

H3K27ac CUT&Tag libraries were prepared as previously described (*11*), with modifications to the protocol. 50,000 cell aliquots were pelleted at 600xg in a swinging bucket rotor centrifuge and washed twice with 200 μL CUT&Tag wash buffer (20 mM HEPES pH 7.5; 150 mM NaCl; 0.5 mM Spermidine (Sigma); 1x Protease inhibitor cocktail (Roche)). Nuclei were isolated by resuspending cell pellets in 200 μL Dig-wash buffer (CUT&Tag wash buffer supplemented with 0.05% digitonin (Promega, G9441) and 0.05% IGEPAL CA-630 (Sigma, I8896)). Nuclei pellets were washed twice with 200 μL of Dig-wash buffer before resuspending in 100 μL Dig-wash buffer supplemented with 2 mM EDTA and a 1:50 dilution of H3K27ac primary antibody (Cell Signaling, 8173) and incubated overnight at 4°C on an overhead rotator. Excess primary antibody was removed by pelleting the nuclei at 600xg and washing twice with 200 μL Dig-wash buffer. Secondary antibody (Novex, A16031) was added at a dilution of 1:50 in 100 μL of Dig-wash buffer, and nuclei were incubated at room temperature for 30 minutes on a rotator. Excess secondary antibody was removed by pelleting the nuclei at 600xg and washing twice with 200 μL of Dig-wash buffer. pA-Tn5 was added at a dilution of 1:100 in 100 μL of Dig-med buffer (0.05% Digitonin, 20 mM HEPES, pH 7.5, 300 mM NaCl, 0.5 mM Spermidine, 1x protease inhibitor cocktail), and nuclei were incubated at room temperature for 1 hour on a rotator. Unbound pA-Tn5 was removed by pelleting the nuclei at 300xg and washing twice with 200 μL Dig-med buffer. Nuclei were resuspended 100 μL of tagmentation buffer (10 mM MgCl_2_ in Dig-med buffer) and incubated for 1 hour at 37°C. After tagmentation, nuclei were lysed with the addition of 100 μL of DNA binding buffer (Zymo Research), and tagmented DNA was purified with a 1.5:1 ratio of AMPure XP beads (Beckman, A63880) following the manufacturer’s instructions. Purified DNA was eluted in 21 μL of EB and mixed with 2 μL each of 10 μM indexed i5 and i7 primers and 25 μL of NEBNext HiFi 2x PCR Master mix. Libraries were amplified with the following conditions: 72 °C for 5 min; 98 °C for 30 s; 12 cycles of 98 °C for 10s and 63 °C for 30 s; final extension at 72 °C for 1 min and hold at 4 °C. Libraries were purified with a 1:1 ratio of AMPure XP beads and eluted in 15 μL of EB. CUT&Tag libraries were quantified by Agilent Bioanalyzer, and sequenced paired-end to a depth of 15 million reads on an Illumina NovaSeq 6000 system, with read lengths 50×8×8×50. Raw basecalls were processed and demultiplexed using Illumina bcl2fastq v2.2, and fastqs were trimmed for the Nextera adapter sequence using TrimGalore v0.6.5, before aligning to the human reference genome (GRCh38) using BWA v0.7.17. pA-Tn5 was provided as a generous gift from Steven Henikoff.

### Enhancer-Gene interaction predictions

Enhancer-gene interactions were predicted using the activity-by-contact algorithm as previously described (*8*). Briefly, a pseudobulk signal from all scATAC microglia cells, bulk H3K27ac CUT&TAG data, publicly available bulk gene expression data (GSM2285362), and an average Hi-C profile from 10 cell lines were used as input to the algorithm. Intersection of the predicted enhancer-gene interactions and cluster-specific peaks was performed using the findOverlaps function from the GenomicRanges R package.

### Primary human microglia purification

Deidentified primary tissue samples were collected with previous patient consent in strict observance of the legal and institutional ethical regulations. Brain tissue was immediately placed in a sterile conical tube filled with oxygenated artificial spinal fluid (aSCF) containing 125 mM NaCl, 2.5 mM KCl, 1mM MgCl_2_, 1 mM CaCl_2_, and 1.25 mM NaH_2_PO4 bubbled with carbogen (95% O_2_/5% CO_2_). Prenatal human microglia were purified from primary brain tissue from mid-gestation (gestational week 18-23) samples using magnetic-activated cell sorting (MACS) kit with CD11b magnetic beads (Miltenyi Biotec, 130-049-601) following manufacturer’s instructions. Briefly, primary brain tissue was minced to 1mm^2^ pieces and enzymatically digested in 10 ml of 0.25% trypsin reconstituted from 2.5% trypsin (Gibco, 15090046) in DPBS (Gibco, 14190250) for 30 mins at 37°C. 0.5 ml of 10 mg/ml of Dnase (Sigma Aldrich, DN25) was added in the last 5 minutes of dissociation. After the enzymatic digestion, tissue was mechanically triturated using a 10 ml pipette, filtered through a 40 μm cell strainer (Corning 352340), pelleted at 300xg for 5 minutes and washed twice with DBPS. Dissociated cells were resuspended in MACS buffer (DPBS with 1 mM EGTA and 0.5% BSA) with addition of 0.5 mg/ml DNAse, and incubated with CD11b antibody for 15 minutes on ice. After the incubation, cells were washed with 10 ml of MACS buffer and loaded on LS columns (Miltenyi Biotec, 130-042-401) on the magnetic stand. Cells were washed 3 times with 3 ml of MACS buffer, then the column was removed from the magnetic field and microglia cells were eluted in 5 ml of MACS buffer. Cells were pelleted at 300xg, re-suspended in 1 ml of culture media and counted and used for downstream analysis. We routinely obtained 1×10^6 of microglia cells per MACS purification.

### Microglia culture

Microglia were cultured on glass-bottom 24 well plates (Cellvis, P24-1.5H-N) pre-coated with 0.1 mg/ml of poly-d-lysine (Sigma Aldrich, P7280) for 1 hr and 1:200 laminin (Thermo Fisher, 23017015) and 1:1,000 fibronectin (Corning, 354008) for 2 hrs. Microglia were plated at 1,5×10^5 cells/well and maintained in culture media containing 66% (vol/vol) Eagle’s basal medium, 25% (vol/vol) HBSS, 2% (vol/vol) B27 (Thermo Fisher, 17504001), 1% N2 supplement (Thermo Fisher, 17502001), 1% penicillin/streptomycin, and GlutaMax (Thermo Fisher) additionally supplemented with 100 ng/ml IL34 (Peprotech, 200-34), 2 ng/ml TGFb2 (Peprotech,100-35B), and 1x CD lipid concentrate (Thermo Fisher, 11905031) for 5-8 days. Media changes were performed twice a week.

### Mouse microglia

Mouse microglia were purified from CD1-IGS (Charles River) P0-P5 mouse pups. Cortices from 4-5 mouse pups were dissected in ice-cold aCSF, combined on a 35 mm dish and minced to 1mm^2^ sections. Cerebral tissue was dissociated and MACS-processed following the same protocol as for the primary human tissue. 1–1.5×10^6 cells were routinely collected for each MACS purification.

### Primary prenatal brain slices

Deidentified primary tissue samples were collected with previous patient consent in strict observance of the legal and institutional ethical regulations. Cortical brain tissue was immediately placed in a sterile conical tube filled with oxygenated artificial spinal fluid (aSCF) containing 125 mM NaCl, 2.5 mM KCl, 1mM MgCl_2_, 1 mM CaCl_2_, and 1.25 mM NaH_2_PO4 bubbled with carbogen (95% O_2_/5% CO_2_). Blood vessels and meninges were removed from the cortical tissue, and then the tissue block was embedded in 3.5% low-melting-point agarose (Thermo Fisher, BP165-25) and sectioned perpendicular to the ventricle to 300 μm using a Leica VT1200S vibrating blade microtome in a sucrose protective aSCF containing 185 mM sucrose, 2.5 mM KCl, 1 mM MgCl_2_, 2 mM CaCl_2_, 1.25 mM NaH_2_PO_4_, 25 mM NaHCO_3_, 25 mM d-(+)-glucose. Slices were transferred to slice culture inserts (Millicell, PICM03050) on six-well culture plates (Corning) and cultured in prenatal brain slice culture medium containing 66% (vol/vol) Eagle’s basal medium, 25% (vol/vol) HBSS, 2% (vol/vol) B27, 1% N2 supplement, 1% penicillin/streptomycin and GlutaMax (Thermo Fisher) additionally supplemented with 100 ng/ml IL34 (Peprotech, 200-34), 2 ng/ml TGFb2 (Peprotech,100-35B), and 1x CD lipid concentrate (Thermo Fisher, 11905031). Slices were cultured in a 37 °C incubator at 5% CO_2_, 8% O_2_ at the liquid-air interface created by the cell-culture insert.

### Slice culture comparison

For the slice culture optimization, we compared slice culture compositions from basal media supplemented with fetal bovine serum (FBS) and in serum-free conditions. Basal media contained 66% (vol/vol) Eagle’s basal medium, 25% (vol/vol) HBSS, 1% N2 supplement, 1% penicillin/streptomycin and 1% GlutaMax (Thermo Fisher). The basal media was used alone or supplemented with 5% (vol/vol) of non-heat-inactivated FBS (nhi-FBS), heat-activated FBS (hi-FBS), or with 2% (vol/vol) B27 with vitamin A. We additionally tested BrainPhys media (BP) (Stemcell Technologies, 05970) supplemented with 1% penicillin/streptomycin,1% GlutaMax, and 1% N2 (BP+N2) or 1% N2+2% B27 (BP+B27+N2).

### Adult human brain slice culture

Adult surgical specimens from epilepsy cases were obtained from the UCSF medical center in collaboration with neurosurgeons with previous patient consent. Surgically excised specimens were immediately placed in a sterile conical tube filled with artificial spinal fluid (aSCF) containing 125 mM NaCl, 2.5 mM KCl, 1mM MgCl_2_, 1 mM CaCl_2_, and 1.25 mM NaH_2_PO_4_ and bubbled with carbogen (95% O_2_/5% CO_2_). The tissue was transported from the operating room to the laboratory for processing within 40–60 min. Blood vessels and meninges were removed from the cortical tissue, and then the tissue block was embedded in 3.5% low-melting-point agarose (Thermo Fisher, BP165-25) and sectioned perpendicular to the cortical plate to 300 μm using a Leica VT1200S vibrating blade microtome in aSCF. Slices were transferred to slice culture inserts (Millicell, PICM03050) on six-well culture plates (Corning) and cultured in adult brain slice culture medium containing 840 mg MEM Eagle medium with Hanks salts and 2mM L-glutamine (Sigma, M4642), 18 mg ascorbic acid (Sigma, A7506), 3 mL HEPES (1M stock) (Sigma, H3537), 1.68 mL NaHCO_3_ (892.75 mM solution, Gibco, 25080-094), 1.126 mL D-glucose, (1.11M solution, Gibco, A24940-01), 0.5 mL penicillin/streptomycin, 0.25 mL GlutaMax (at 400x, Gibco, 35050-061), 100 μL 2M stock MgSO_4_.7H_2_O (Sigma, M1880), 50 μL 2M stock CaCl_2_.2H_2_O (Sigma, C7902), 50 μL insulin from bovine pancreas, (10 mg/mL, Sigma, I0516), 20 mL horse serum-heat inactivated, 95 mL MilliQ H_2_O (as previously described (*12*)). Slices were cultured at the liquid-air interface created by the cell-culture insert in a 37°>C incubator at 5% CO_2_ for two days before primary prenatal microglia transplantation.

Primary prenatal human microglia were MACS-purified as described above and labeled with DiI (Thermo Fisher, V22885) following manufacturer’s instructions. Briefly, 5 μL of the cell labeling DiI solution was added to 1 mL of microglia cell suspension. Cells were incubated at 37°C for 10 minutes and washed twice with DPBS. 1,5×10^5 cells were added to each adult brain slice and cultured for five additional days.

### Organoid generation

Cerebral organoids were generated based on a previously published method (*13*) with several modifications. Briefly, hiPSCs cultured on Matrigel were dissociated into clumps using 0.5 mM EDTA in Ca^2+^/Mg^2+^-free DPBS and transferred into ultra-low attachment 6-well plates in neural induction media (GMEM containing 20% (v/v) KSR, 1% (v/v) penicillin-streptomycin, 1% (v/v) non-essential amino acids, 1% (v/v) sodium pyruvate, and 0.1 mM 2-mercaptoethanol). For the first nine days, neural induction media was supplemented with the SMAD inhibitors SB431542 (5 μM) and dorsomorphin (2 μM) and the Wnt inhibitor IWR1-ε (3 μM), with a media exchange performed every three days. Additionally, the Rho Kinase Inhibitor Y-27632 (20 μM) was added during the first six days of neural induction to promote survival. Between days 9-25, organoids were transferred to a neural differentiation media (1:1 mixture of Neurobasal and DMEM/F12 containing 2% (v/v) B27 without vitamin A, 1% N2, 1% (v/v) non-essential amino acids, 1% (v/v) Glutamax, 1% (v/v) antibiotic/antimycotic, 0.1 mM 2-mercaptoethanol) supplemented with FGF2 (10 ng/mL) and EGF (10 ng/mL). Between days 25-35, organoids were maintained in neural differentiation media without FGF or EGF. From Day 35 onward, organoids were maintained on a shaker in neural differentiation media containing B27 with vitamin A with media exchanges every 2-3 days.

### Microglia transplantation

MACS-purified microglia were added to week 5 organoids in 6-well plates at 1×10^5 microglia cells/organoid and kept off the shaker overnight. The next day, the plates were returned to the shaker and maintained following the usual organoid maintenance protocol.

### Organoid single-cell capture for single-cell RNA sequencing

Two organoids per experimental condition were cut into 1 mm^2^ pieces and enzymatically digested with pa apain digestion kit (Worthington, LK003163) with the addition of 0.5 mg/ml DNase for 1 hr at 37°C. Following enzymatic digestion, organoids were mechanically triturated using a P1000 pipette, filtered through a 40 μm(*14*)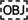 for multiplexing. Three organoid lines with and without microglia were combined and captured on two lanes of 10x Genomics using Chromium single cell 3’ reagent kit (v3 Chemistry) following the manufacturer’s protocol.

### Antibodies

Primary antibodies used in this study included: rabbit Iba1 (1:500, Wako, 019-19741), guinea pig Iba1 (1:500, Synaptic Systems, 234 004), mouse VGLUT1 (1:200, Millipore Sigma MAB5502), rabbit psd95 (1:350, Thermo Fisher, 51-6900), rabbit P2RY12 (Sigma, HPA014518). Secondary antibodies were species-specific AlexaFluor secondary antibodies (1:1,000)

### Immunofluorescence of cryosections

Organoids and primary human brain tissue samples were fixed in 4% PFA for 1 hour. Tissue sections were cryopreserved in OCT/30% sucrose (1:1) and cryosectioned at 20 μm or 40 μm (for synaptic density staining) thickness. Heat-induced antigen retrieval was performed in 10mM sodium citrate (pH=6.0) for 10 min in boiling-hot solution. Blocking and permeabilization were performed in a blocking solution consisting of 10% normal donkey serum, 1% Triton X-100, and 0.2% gelatin for 1 hour. Primary and secondary antibodies were diluted and incubated in the blocking solution. Cryosections were incubated with primary antibodies at 4^0^C overnight, washed 3x with washing buffer (0.1% Triton X-100 in PBS). Sections were incubated with secondary antibodies in the blocking buffer for 1 hour at the room temperature. Images were collected using Leica SP8 confocal system with 20x air objective and 63x oil objective and processed using ImageJ/Fiji and Illustrator.

### Immunofluorescence of adult human brain slices

Adult human brain slices were fixed in 4% PFA for 1 hour. The immunofluorescence staining was performed as for the cryosections with the difference that Triton-X was omitted from the blocking buffer. Heat-induced antigen retrieval was performed in 10mM sodium citrate (pH=6.0) for 10 min in boiling-hot solution. Blocking and permeabilization were performed in a blocking solution consisting of 10% normal donkey serum, 0.5% saponin, and 0.2% gelatin for 1 hour. Primary and secondary antibodies were diluted and incubated in the blocking solution. Brain slices were incubated with primary antibodies at 4^0^C overnight and washed 3x with PBS followed by incubation with secondary antibodies in the blocking buffer for 1 hour at the room temperature. Images were collected using Leica SP8 confocal system with 20x air objective and 63x oil objective and processed using ImageJ/Fiji and Illustrator.

### Immunofluorescence of 2D cell cultures

Cells cultured on glass-bottom well plates were fixed in 4% PFA for 10 minutes. Blocking and permeabilization were performed in a blocking solution consisting of 10% normal donkey serum, 1% Triton X-100, and 0.2% gelatin for 1 hour. Primary and secondary antibodies were diluted and incubated in the blocking solution. Cell cultures were incubated with primary antibodies at the room temperature for 1 hour, washed 3x with washing buffer (0.1% Triton X-100 in PBS), and incubated with secondary antibodies for 1 hour at the room temperature. Images were collected using Leica SP8 confocal system with 20x air objective and 63x oil objective and processed using ImageJ/Fiji and Illustrator.

### Synapse quantification

Organoid confocal images were acquired with a Leica SP8 system. For synapse quantification, three different optical fields per organoid were imaged. For each optical field, 15 μm Z-stack (0.5μm Z-step) were collected using a 63X/1.40 oil objective. Synapse determination was based on the colocalization between Vglut1 (1:200; Millipore Sigma MAB5502) and PSD95 (1:350; Invitrogen 51-6900) as previously described (*15*) using ImageJ software. Briefly, the background was subtracted with a rolling bar radius of 10 pixels. Then, threshold was applied to every channel in order to distinguish synaptic puncta from background and generate two new binary images with the synaptic markers. Colocalization was determined overlaying both binary images with the synaptic markers. Finally, synaptic puncta were determined using the function “analyze particles’’.

### Single-molecule RNA in situ hybridization

RNA *in situ* hybridization experiments were performed using the RNAscope^®^ technology, which has been previously described (*16*). Paired double-Z oligonucleotide probes were designed against target RNA using custom software. The following probes were used: *Hs-HSPA6-O1-T1*, Hs-*P2RY12*-T2, Hs-*IL1B*-T3, Hs-*CCL2*-T4, Hs-*APOE*-T5, Hs-*CX3CR1*-T6, Hs-*LGALS3*-T7, Hs-*MIF*-T8. The RNAscope HiPLex8 reagent kit (Advanced Cell Diagnostics, 324100) was used according to the manufacturer’s instructions. PFA-fixed frozen tissue sections were processed according to manufacturer’s recommendations. Sagittal sections from two individuals, GW20 and GW13 were used, with three imaging fields per anatomical region (thalamus and across the cortex, spanning ventricular zone and cortical plate) collected from each sample. Fluorescence images were acquired using a Leica SP8 microscope using a 20x objective. Probes were considered as positive when more than 3 puncta overlapping with DAPI staining were detected. To limit the analysis to RNA expression in microglia cells exclusively, only probes that overlayed with *P2RY12* RNA probes were included in the analysis. Individual image stacks were registered, cropped and aligned using the ACD HiPlex software package for Mac OS. Microglia cell identity was additionally confirmed with the *CX3CR1* probe, but only *P2RY12* was used for quantification. The same tissue sections used for hybridization were used for immunofluorescence analysis to visualize cell outlines and were combined with the RNAscope data using the HiPlex software.

### Axion recordings

Multielectrode array recordings were conducted using the Axion Maestro Pro system 64 low-impedance PEDOT electrodes with 300 μm electrode spacing. Briefly, organoids were transferred to a 64-channel multielectrode plate and equilibrated at 37°C, 5% CO_2_ for at least 15 minutes prior to recording. Spontaneous neural activity was recorded for 40 mins. Firing events were defined as events with threshold amplitude of more than six standard deviations of the background noise. The event amplitude ranged between 8 μV to over 60 μV for some events. The majority of events consisted of multi-unit events and demonstrated waveforms characteristic for a firing neuron. The recordings at 12,500 samples/second were performed in the same media used for organoid growth. After recording, organoids were detached from the MEA plate and kept on the shaker.

### Axion data analysis

Raw data from the Axion recordings were read into Matlab then converted to numpy arrays for further processing in Python. Spikes were counted using a standard deviation threshold of 6 and converted to an array of spike times for each individual channel. Bursting activity was determined by the presence of 10 or more spikes on a given channel within one second. Next, recording time was divided into 40 ms bins and spike events were binarized based on whether or not there was at least one spike within a given time bin. Pairwise correlation between channels was determined by calculating the Pearson correlation between these binarized spike times.

